# Retinal OFF ganglion cells allow detection of quantal shadows at starlight

**DOI:** 10.1101/2021.11.02.466884

**Authors:** Johan Westö, Nataliia Martyniuk, Sanna Koskela, Tuomas Turunen, Santtu Pentikäinen, Petri Ala-Laurila

## Abstract

Perception of light in darkness requires no more than a handful of photons and this remarkable behavioral performance can be directly linked to a particular retinal circuit – the retinal ON pathway. However, the neural limits to shadow detection in very dim light have remained unresolved. Here, we unravel the neural mechanisms that determine the sensitivity of mice to the dimmest light decrements by measuring signals from the most sensitive ON and OFF retinal ganglion cell types and by correlating their signals with visually guided behavior. We show that mice can detect shadows when only a few photon absorptions are missing among thousands of rods. Behavioral detection of “quantal” shadows relies only on the retinal OFF pathway and is limited by noise and losses of single-photon signals in retinal processing. Thus, in the dim-light regime, light increments and decrements are encoded separately via the ON and OFF retinal pathways, respectively.

## INTRODUCTION

Dark-adapted humans can detect weak flashes of light leading to a dozen or so absorbed photons in the retina. This outstanding sensitivity for light increments in darkness approaches the ultimate limits set by the quantal nature of light, the neural noise, and the unavoidable losses of single photons in their capture and processing in the retina (reviewed in Kiani et al., 2020; Schwartz, 2021). Such an exquisite sensitivity requires a set of well-optimized neural mechanisms end-to-end from the retina to visually-guided behavior. However, behavioral and neural limits for detecting the weakest light decrements – quantal shadows – have hardly been addressed. This is surprising given the obvious functional importance of detecting dark prey and predators as decrements or shadows even in starlight. Indeed, psychophysical studies on humans have established that the detection thresholds in terms of light quanta are lower for decrement than for increment contrast stimuli at scotopic light levels (Blackwell, 1946; Patel and Jones, 1968). Similarly, behavioral studies on mice have shown that decrements can already be discerned at starlight levels (Sampath et al., 2005). However, direct links between behavioral sensitivity, the underlying retinal output signals provided by retinal ganglion cells (RGCs), and the theoretical limits set by photon distributions have not been established for light decrements.

In starlight, only a few rods among approximately a thousand absorb photons, whereas the rest of the rods generate noise. Thus, the visual world is no longer continuous, and even shadows correspond to a small number of “missing” photons in the sprinkle of sparse photons originating from the background light (see Figure 1). In these dimmest conditions, the mammalian vision relies on a well-known retinal circuit, the rod bipolar pathway (Bloomfield and Dacheux, 2001; reviewed in Field et al., 2005). The sparseness of visual signals and the rich knowledge of neural processing principles along this pathway offer a unique possibility for comparing retinal signals directly to the theoretical limits arising from the photon distribution. Two key features of the rod bipolar pathway are: 1) A continuous increase in the level of rod convergence, with cells at later stages pooling input from increasing numbers of rods and thereby allowing these later stages to better discriminate stimulus-elicited responses from the background light. For example, single rods are unable to statistically discern a quantal shadow, as they rarely even absorb a single photon at these light levels, whereas the most sensitive RGCs pool inputs from thousands of rods, allowing a much better discrimination of sparse stimuli (see Figure 1). 2) The circuit is split into separate ON and OFF pathways at a late stage (at the level of the AII amacrine cells). ON RGCs respond to light increments with increasing firing rates and light decrements with decreased firing rates, whereas OFF RGCs have the opposite response polarities. ON and OFF RGCs thus share much of the same well-known circuitry and pooling mechanisms of the single-photon responses, with pathway-specific differences introduced in the last synapse (Ala-Laurila and Rieke, 2014).

**Figure 1.**
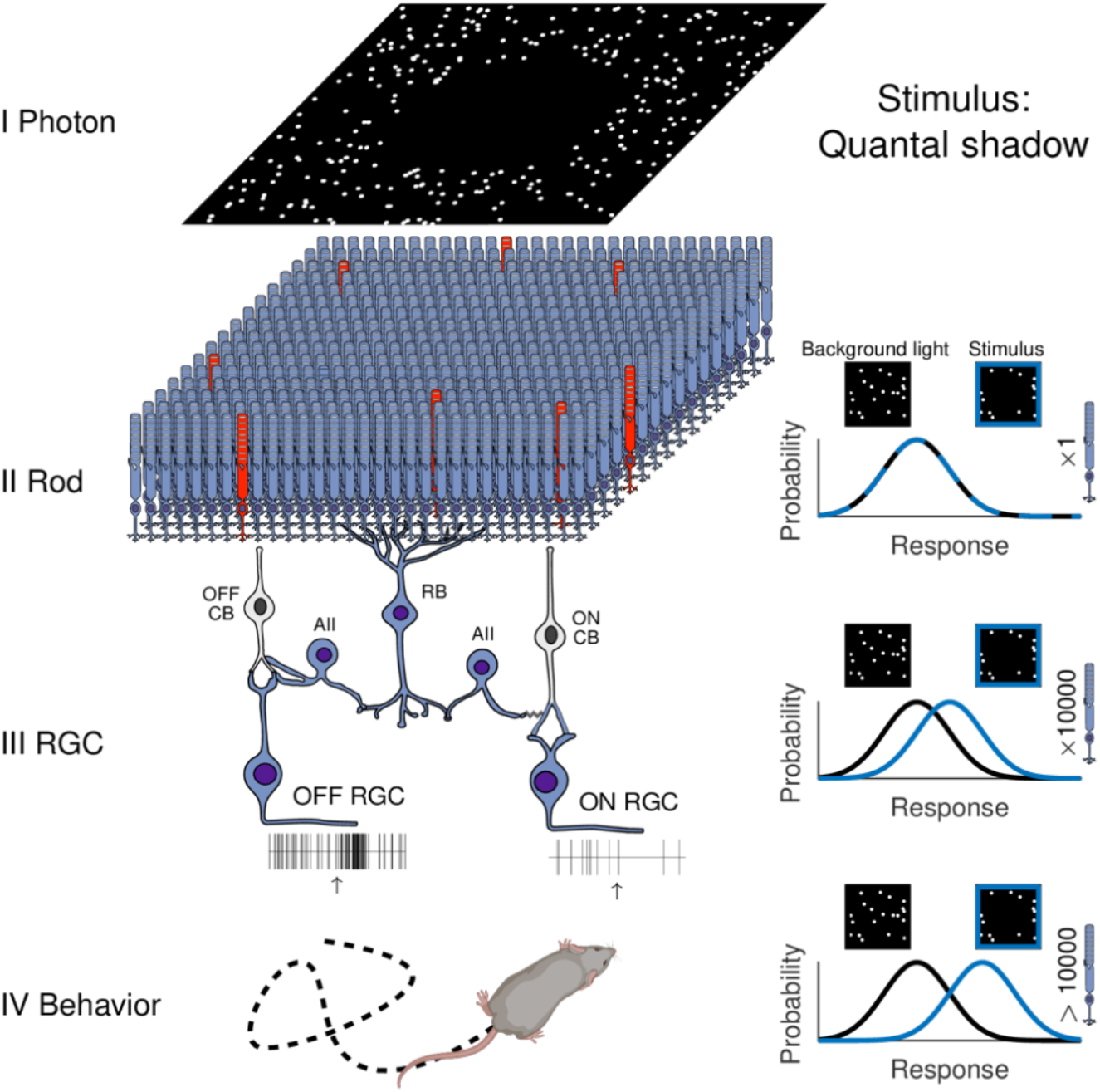
Decrement detection at very dim background light requires photon responses to be pooled from thousands of rods. In starlight conditions, light decrements correspond to missing photons in a sparse photon sprinkle (“quantal shadows”). Only a small proportion of all photons incident on the eye are absorbed in rods, which makes the quantal shadows even more difficult to detect in the rod array, where only a few rods (red) among thousands may exhibit a single-photon response. The curves in the right-hand panels illustrate the probability distributions of responses to background light (black) and to quantal shadow stimuli (blue) at three primary levels of the visual system (Rods, RGCs, and visually guided behavior). Note that a single rod can never distinguish the lack of photons due to a quantal shadow on its own (the distributions coincide), as a single rod hardly ever absorbs a photon in very dim backgrounds. The mammalian retina therefore routes single-photon responses through a specialized neural circuit, the rod bipolar pathway (shown on the left, blue color). This pathway is characterized by a successively increasing rod convergence as the signal traverses the rod-bipolar cells (RB), the AII amacrine cells, and finally ON and OFF alpha RGCs (see rod convergence numbers, bottom right). Each RGC pools input from ~10,000 rods, thus enabling detection of very faint light decrements that would be impossible to statistically discern in single rods, or in a small number of rods. At the level of behavior, the brain can further pool over multiple RGCs to detect decrements in even dimmer backgrounds. The increasing distance between the background-light-elicited distribution (black) and signal-elicited response distributions (blue) through levels II–IV shows the impact of the increased rod pooling across the rod bipolar pathway. OFF CB = OFF cone bipolar cell, ON CB = ON cone bipolar cell.

Previous electrophysiological studies from the lateral geniculate nucleus (LGN) and behavioral studies using visually guided saccadic reaction times in monkeys have shown that pharmacological blockage of the ON pathway compromises detection of light increments, but not that of light decrements, at cone-driven light levels (Schiller et al., 1986; Schiller, 2010). Similarly, theoretical arguments suggest that the split of visual information into parallel ON and OFF pathways allows overall more efficient coding of visual scenes than a single pathway (Gjorgjieva et al., 2014). However, two major technical challenges have made the direct linkage between retinal outputs and behavioral performance impossible. First, both ON and OFF pathways can encode both positive and negative contrast at photopic light levels with increasing and decreasing firing rates to light increments and decrements, and their precise potential for mediating this information depends strongly on stimulus conditions (Liang and Freed, 2010; Manookin et al., 2008). Thus, the linkage between visually guided behavior and ON and OFF RGC responses requires RGC recordings and behavioral measurements in precisely matching conditions. Second, photopic vision relies on several tens of distinct RGC types in the mammalian retina (Baden et al., 2016; Bae et al., 2018; Rheaume et al., 2018; Tran et al., 2019; Field and Chichilnisky, 2007; Peng et al., 2019) making it extremely difficult to link RGC-type-specific retinal outputs to behavior. Starlight, in contrast, offers a unique possibility for this (reviewed in Kiani et al., 2020), as the relevant retinal output is constrained to the most sensitive RGC types. Relying on this key insight, we have previously linked the most sensitive ON RGCs to increment detection in darkness in both mice (Smeds et al., 2019) and primates, including humans (Kilpeläinen et al., 2021). However, the behavioral relevance of OFF RGCs in scotopic conditions has remained unresolved.

Here, we have recorded decrement responses of the most sensitive ON and OFF RGCs in mice (alpha RGCs) and evaluated their ability to encode negative contrast in very dim background lights. We compared the spike responses of these RGCs with 1) measured behavioral performance on a decrement detection task near the sensitivity limit of vision, and 2) ultimate theoretical limits for decrement detection as estimated from photon distribution at different levels of the retinal circuit mediating these signals. We found that behavior reflects a near-optimal readout of the OFF RGC spike code and that the deviation from this limit is almost exclusively due to losses of single photons and single-photon responses as well as neural noise in the retina.

## RESULTS

Our first goal was to characterize the spike responses of the most sensitive mouse RGC types to extremely dim decrement light stimuli, hereafter referred to as quantal shadows. We have previously identified the three alpha RGC types in the mouse retina (OFF sustained, OFF transient, and ON sustained, hereafter referred to as OFF-S, OFF-T, and ON-S) as the most sensitive cell types for light increments in darkness, where RGCs receive their inputs via the rod bipolar pathway (Koskela et al., 2020; Smeds et al., 2019). Here, we recorded their spike responses both to light increments in darkness and to light decrements across a range of dim background lights eliciting 0.03–30 photoisomerizations per rod per second (R*/rod/s). The lower end of this range (< 1 R*/rod/s) is encoded solely by the rod bipolar pathway (Grimes et al., 2018). We focused on the two sustained alpha types, OFF-S and ON-S, as a proxy for the most sensitive readout of the retinal ON and OFF pathways. Although OFF-S and OFF-T RGCs have very similar sensitivities for flashes of light in darkness (Murphy and Rieke, 2011), the long stimulus durations needed for detecting quantal shadows favor OFF-S RGCs, thus making them the best proxy for the most sensitive readout of the OFF pathway.

The stimulus space for quantal shadows is fundamentally different from that for light increments in darkness, as demonstrated in Figures 2A and 2B. The key difference is that the background light level sets an upper limit for the maximal stimulus magnitude for light decrements (but not for light increments) of any specific duration. For example, a 20 ms decrement stimulus of maximal contrast will, on average, remove 40 R* per RGC receptive field under a background of 0.2 R*/rod/s, but only 2 R* under a background of R*/rod/s (assuming a rod convergence of 10 000 per RGC; Dunn et al., 2006; Tsukamoto et al., 2001). The latter is well below the detection threshold of even the most sensitive RGCs in mice (Koskela et al., 2020; Takeshita et al., 2017). As the contrast cannot be increased beyond 100%, it is possible only by prolonging the stimulus duration to remove 40 R* at 0.01 R*/rod/s (Figure 2B). Thus, increasing the stimulus duration is the only means of increasing the stimulus magnitude for the dimmest quantal shadows. Figure 2B highlights this effect by showing stimulus magnitudes for stimulus durations up to 400 ms, corresponding approximately to the rod integration time (~370 ms; Azevedo and Rieke, 2011), which is the longest known integration time across the cell types in the rod bipolar pathway.

**Figure 2.**
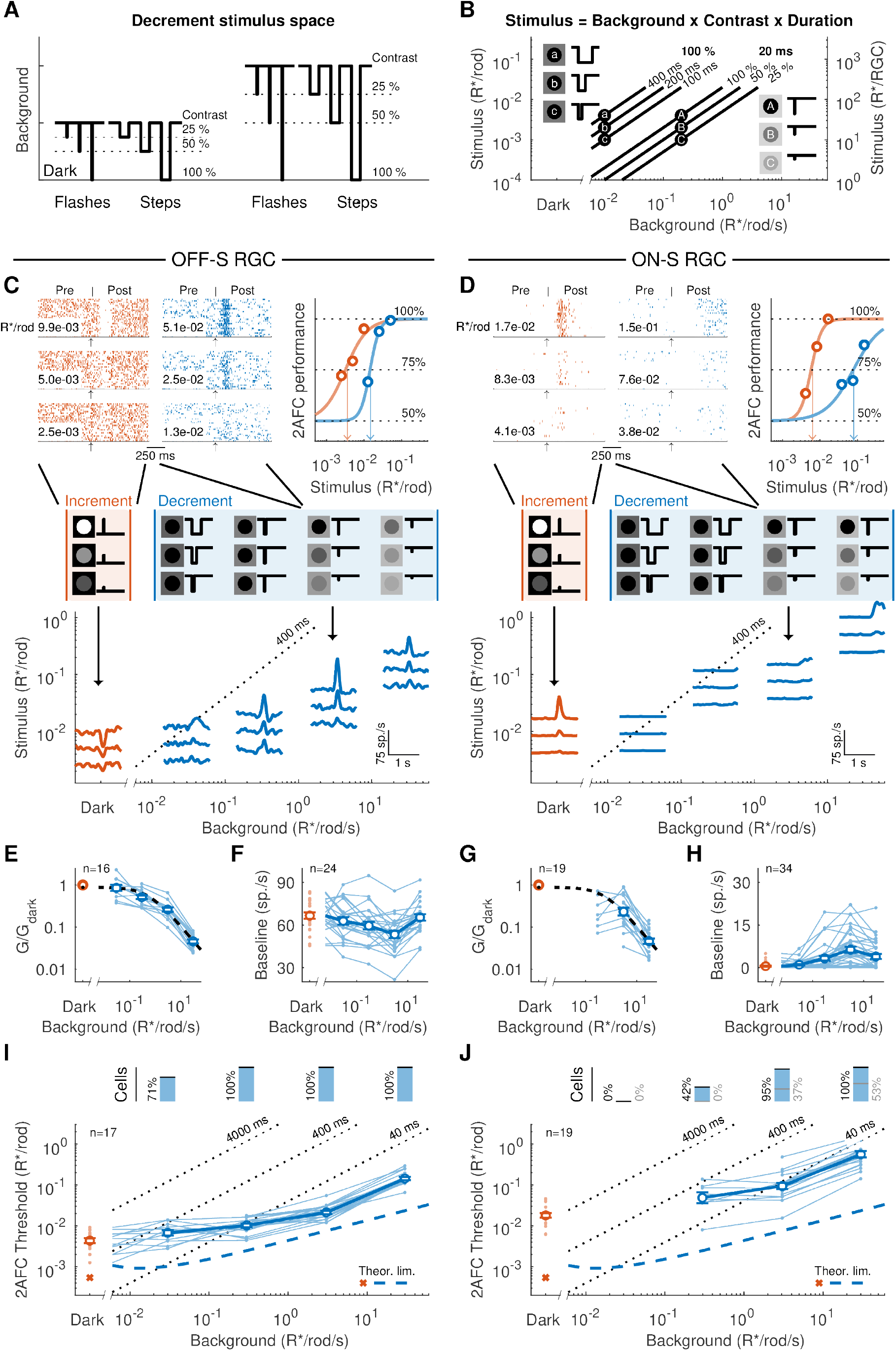
Comparison of OFF-S and ON-S RGC responses to light increments in darkness and light decrements across dim backgrounds. (A) The absolute magnitude of flash and step decrement stimuli of fixed contrast and duration scale with background intensity. (B) Brief decrement flashes fail to elicit RGC responses below a certain background intensity, which means that the duration of the decrement needs to be prolonged in order to present stimuli of matching magnitude when lowering the background intensity (A to a, B to b, and C to c). The black lines depict how the magnitude of each stimulus changes as a function of the background. (C) Peri-stimulus time histograms (bottom) for one OFF-S RGC in response to increment (red) flashes of three magnitudes in darkness, and in response to decrements (blue) of three magnitudes at four background levels (0.03, 0.3, 3, and 30 R*/rod/s). Spike rasters (top left) are shown for the increment responses in darkness and for the decrement responses at 3 R*/rod/s. The weakest stimulus encoded in the spike response was quantified by evaluating how frequently an ideal observer could determine whether the time interval preceding the stimulus (Pre) or the time interval after the stimulus (Post) of the spike train contained the response (top right). (D) Same as (C), but for an ON-S RGC. (E) Sensitivities (spikes/R*) for OFF-S RGCs in response to decrements at all four background intensities (normalized by the sensitivity to increments in darkness). The dashed line denotes a fitted generalized Weber function (Equation 4) with the parameters G_D_ = 0.9, I_o_ = 0.6, and m = −0.7. (F) Baseline firing rates of OFF-S RGCs in darkness and at the four tested backgrounds. (G) and (H) Same as (E) and (F), but for ON-S RGCs. The observed spontaneous firing rates in darkness (red symbols in Figures 2H & 2F) agreed with earlier studies (Koskela et al., 2020; Takeshita et al., 2017). (I) Two-alternative forced-choice (2AFC) thresholds for OFF-S RGCs in response to increments in darkness and to decrements at the four background intensities. The bars above represent the fraction of cells encoding a decrement stimulus at each background. The dotted black lines denote the maximal decrement possible for various stimulus durations. (J) Same as (I), but for ON-S RGCs with the gray numbers further indicating the fraction of ON cells that encode decrements through increased firing to the light increase at the cessation of the decrement stimuli. All error bars denote SEMs.

We first recorded RGC responses to brief flashes of light in darkness (red post-stimulus-time histograms (PSTHs), and rasters in Figures 2C and 2D) to validate the sensitivity of all preparations, and we only included preparations with similar sensitivity as in previously published studies (see STAR Methods and Koskela et al., 2020; Smeds et al., 2019; Takeshita et al., 2017). We then recorded the responses of OFF-S and ON-S RGCs to the novel quantal shadow stimuli of three different stimulus magnitudes (blue PSTHs and rasters in Figures 2C and 2D) at four different background light intensities (0.03, 0.3, 3, and 30 R*/rod/s). The ability of OFF-S and ON-S RGCs to encode quantal shadows differed considerably. OFF-S RGCs responded with increasingly higher firing rates as the stimulus magnitude grew. Their sensitivity (spikes/R* = response gain, G) to increments from darkness and decrements in background lights was nearly identical up to backgrounds of ~0.3 R*/rod/s, whereupon light adaptation effects in response gain became prominent (Figure 2E). The overall background dependence of OFF-S RGC decrement responses could be well described with a generalized Weber function with a slope of −0.7 (Figure 2E), in line with previous findings using light increments across dim backgrounds and ON-S RGCs (Dunn et al., 2006). ON-S RGCs, in contrast, responded to decrements of increasing magnitude by progressively decreasing their firing rate (Figure 2D). However, the low baseline firing rate limited the possibility of ON-S RGCs to encode light decrements at these dim background light levels. Only decrements strong enough to elicit a rebound response of opposite polarity at stimulus offset were associated with significant changes in spike counts (Figure 2D). However, no rebound responses were elicited under the dimmest background lights tested (0.03 R*/rod/s), not even with the longest decrement pulses (400 ms) tested. Consequently, ON-S RGCs did not change their firing rate at all in response to quantal shadows at the dimmest background lights tested (Figures 2D and 2G).

Under brighter backgrounds (> 1 R*/rod/s), the baseline firing rates of ON-S RGCs increased, and the cells became correspondingly more responsive to quantal shadows (Figure 2H). Under these background lights, they also exhibited similar light adaptation as the OFF-S RGCs (Figure 2G). Interestingly, the changes in baseline firing rate in ON-S and OFF-S RGCs across this intensity range (darkness to 30 R*/rod/s) were like mirror images: that of ON-S RGCs grew gently with increasing background intensities, up to a maximum of 6 ± 1 spikes/s (mean ± SEM) at 3 R*/rod/s (blue symbols in Figure 2H), whereas that of the OFF-S RGCs decreased gently to reach a minimum at the same background intensity (Figure 2F). The mirrored change in baseline firing rates between ON-S and OFF-S RGCs across dim background lights is in line with the known shared circuitry along the rod bipolar pathway as far as the AII amacrine cells, which provides an excitatory drive to ON-S via ON cone bipolar cells and a direct inhibition to OFF-S. These opposing effects of AII amacrine cells to the ON and OFF pathways supposedly cause the negative correlation of ON-S and OFF-S RGC firing rates across dim backgrounds (see also Murphy and Rieke, 2006).

Next, we addressed our primary question at the level of RGCs: what are the dimmest quantal shadows that the most sensitive ON and OFF RGCs can encode with their spike output? We quantified the ability of ON-S and OFF-S RGCs to encode light increments (in darkness) and decrements in dim backgrounds with an ideal observer analysis considering both the stimulus-elicited responses and the tonic firing rates (Ala-Laurila and Rieke, 2014; Chichilnisky and Rieke, 2005; Smeds et al., 2019). The increment thresholds (defined as 75% correct choices in a two-alternative forced-choice task) were slightly lower for OFF-S RGCs as compared to ON-S RGCs (p = 1.2e-5, Welch’s t-test), and in agreement with previous results (ON & OFF parasols in primate: Ala-Laurila and Rieke, 2014; ON-S and OFF-S in mice: Koskela et al., 2020; Smeds et al., 2019). However, the thresholds for quantal shadows differed fundamentally between ON-S and OFF-S RGCs. OFF-S RGCs encoded decrements down to the dimmest backgrounds tested and reached almost equally low thresholds for quantal shadows and light increments in darkness (Figure 2I). ON-S RGCs, in contrast, did not respond to quantal shadows at all at the dimmest backgrounds (< 0.3 R*/rod/s). ON-S RGCs encoded decrements reliably only under the two brightest backgrounds (3 and 30 R*/rod/s, see Figure 2J), and even then, roughly half of the ON-S RGCs did so by increasing their firing rates in response to the light increase at the cessation of the decrement stimuli (Figure 2J, gray numbers). These results demonstrate that OFF-S RGCs carry reliable information about quantal shadows in dim backgrounds, whereas ON-S RGCs do not.

### Mice can see quantal shadows corresponding to a few missing photons in thousands of rods

Now that we have observed a sensitivity asymmetry between ON-S and OFF-S RGC outputs for quantal shadows, we used this asymmetry to address our primary question: How does behavioral detection of quantal shadows (negative contrast at the lowest light levels) depend on retinal ON and OFF pathways? We established a behavioral paradigm allowing us to measure the behavioral detection limit under stimulus conditions closely matching those in the RGC measurements (Figure 3A). Mice had to detect a black stimulus spot that signaled the presence of a transparent escape ramp from the water in one out of six corridors in a dimly illuminated six-armed white water maze. The swimming behavior was recorded using our markerless, fully automated tracking of mouse head location and direction (Figure 3B and Video S1; Koskela et al., 2020; Smeds et al., 2019). We quantified the behavioral detection limit by measuring the frequency of finding the stimulus corridor as a function of the background intensity.

**Figure 3.**
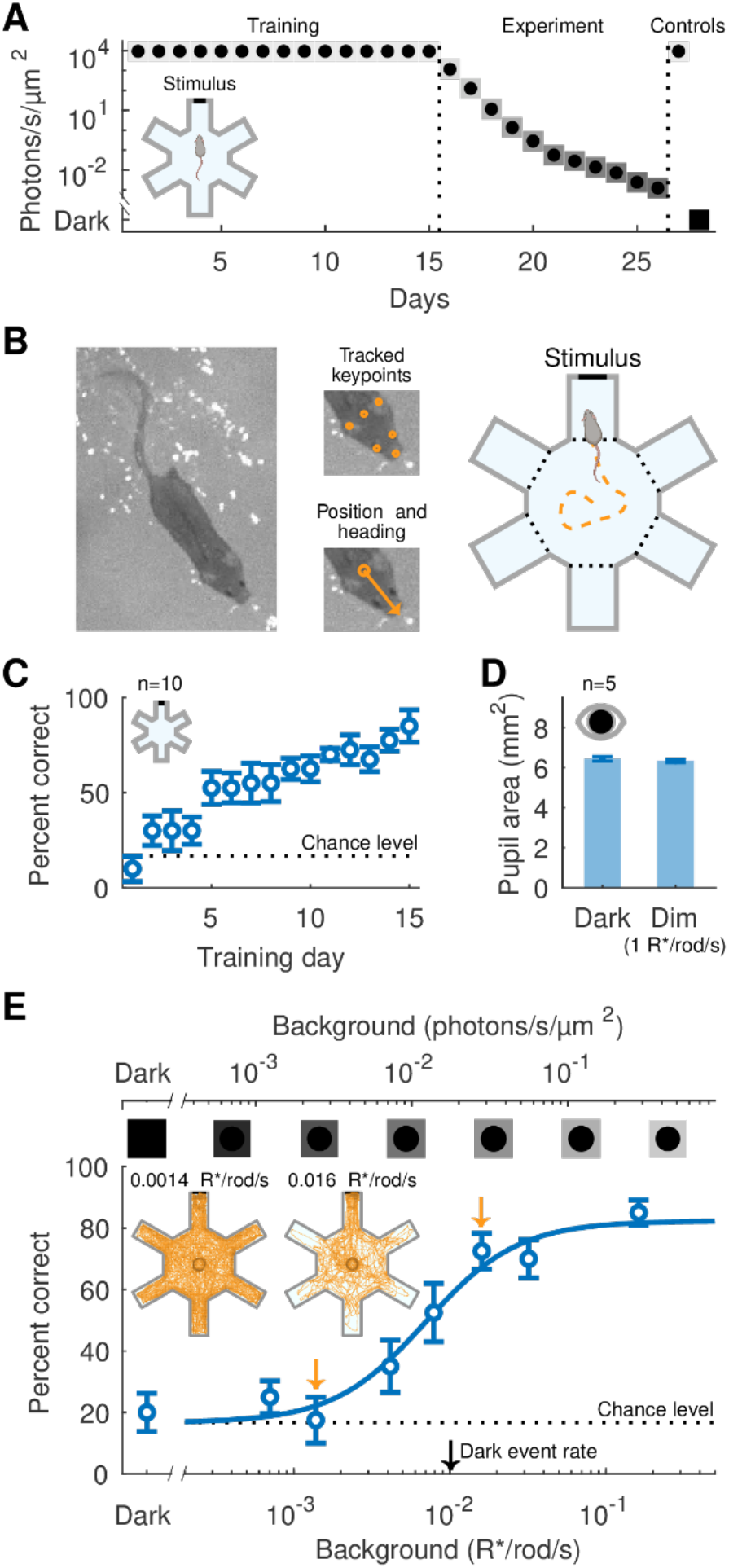
Evaluation of the sensitivity limit for behavioral performance on a decrement detection task. (A) The experimental protocol used to evaluate the sensitivity limit of decrement detection using a homogeneously lit six-armed white water maze (inset). The protocol consisted of a training phase with a fixed background (15 days), an experiment phase during which the background was successively attenuated (11 days), and a control phase where the mice were re-tested with the training background and in complete darkness (2 days). (B) The mice were filmed, and a deep neural network extracted 6 keypoints, which were fused to position and direction estimates and finally filtered to track complete swimming paths. (C) Performance was evaluated using the fraction of correct choices, with the choice defined as the first corridor entered by the mouse. Performance successively improved during training and surpassed our learning criterion of 80% after 15 days (the dashed line indicates the chance level ~17%). Each data point represents the mean over four trials for every mouse (n = 10). (D) Pupil areas measured in darkness and at 1 R*/rod/s (n = 5 mice). (E) Behavioral performance on the decrement detection task (n = 10 mice). The smooth fit is given by Equation 5 with the parameters *K* = 0.007 and *n* = 1.5. The insets (top left) show heat maps for tracked swimming paths for 40 trials at the two intensities highlighted by the orange arrows (0.0014 R*/rod/s and 0.016 R*/rod/s). The black arrow denotes the dark event rate in rod photoreceptors (thermal isomerizations; Burns et al., 2002), and the horizontal dotted line denotes the chance level (~17%). All error bars represent SEMs.

The experimental protocol (Figure 3A) followed a similar structure as previously used for light increment detection (Smeds et al., 2019), with a training phase at constant background intensity followed by the experiments with successively decreasing background intensities. The mice learned the task in about two weeks (surpassing 80% correct choices after 15 days; Figure 3C), in agreement with previous results on light increment detection (Koskela et al., 2020; Smeds et al., 2019). In the subsequent experimental phase, the frequency of finding the stimulus was measured across a number of background light intensities in decreasing order, testing one background light intensity per day. Pupil measurements allowed us to convert all light intensities to retinal isomerization rates (see STAR Methods and Smeds et al., 2019). There was no difference in pupil area between darkness and the background light of 1 R*/rod/s (Figure 3D; p = 0.56, paired t-test), consistent with previous results at similar light levels (Sampath et al., 2005) and the idea that the sustained component of the pupillary light response is not modulated at the dim light levels where signals are mediated by the primary rod pathway (Grimes et al., 2018).

Behavioral detection of quantal shadows showed a monotonic improvement in performance over the range 0.0014–0.016 R*/rod/s, from chance level (dashed line), i.e. 1/6 = 17% correct, to steady-state high performance, i.e. > 80% correct (Figure 3E). The insets in Figure 3E illustrate population swimming trajectories recorded at both ends of this range and highlight how the mice successively transition from a random exploration of all corridors to almost exclusively entering the stimulus corridor at backgrounds just below R*/rod/s. A comparison of this behavioral performance to individual ON-S and OFF-S sensitivities highlights that behavioral performance saturates (> 80% correct) at backgrounds ~10-fold dimmer than the sensitivity limit for the most sensitive ON-S RGCs in encoding quantal shadows. OFF-S RGCs, in contrast, did respond to quantal shadows all the way down to these dim background lights relevant for behavioral detection threshold. Surprisingly, the mice detected quantal shadows in background lights dimmer than the spontaneous isomerization rate of rhodopsin (pigment noise) in single rods (0.01 R*/rod/s; Burns et al., 2002; Fu et al., 2008), indicating that behavioral detection must rely on a considerable amount of pooling of rod signals in both space and time.

### Behavioral performance is in line with a near-perfect readout of the most sensitive OFF RGCs

The similarity between behavioral performance and the performance of single OFF-S RGCs, but not ON-S RGCs, suggests that behavior relies on the retinal OFF-pathway. However, the comparison is complicated by the fact that behavior can pool sparse signals from neighboring RGCs as the small stimulus spot moves over the mosaic of RGCs (see Figures 4A and 4B). To link performance between RGCs and behavior, we thus took a similar approach as in our earlier work for light increments (Smeds et al., 2019). The logic of this approach is as follows (see Figures 4A–C): 1) The tracked position and direction of the mouse head lets us calculate the projection of the stimulus spot on retinal mosaics of ON-S and OFF-S RGCs across the entire swimming trajectory during our behavioral experiments. 2) We simulated ON-S and OFF-S RGC spike trains from the full mosaics by relying on models constrained by RGC spike recordings in matching stimulus conditions (Figures 2C and 2D). 3) We fed ON-S and OFF-S RGC population responses from the full mosaics across the swimming trajectory to ideal observer models (one model relying on the OFF-S RGC population responses – the OFF model – and the other model on the ON-S RGC population responses – the ON model) to evaluate ideal performance on the quantal shadow detection task.

**Figure 4.**
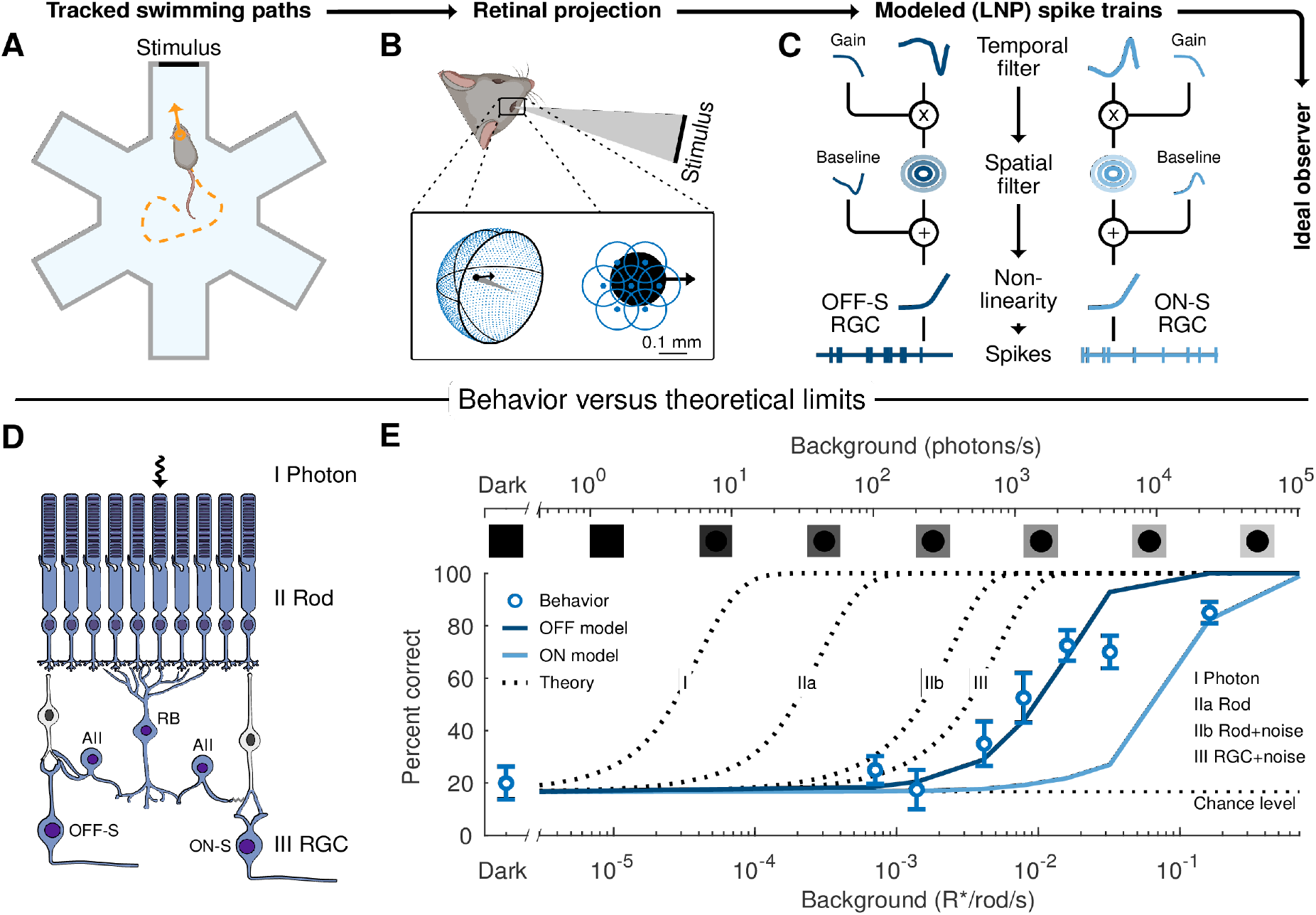
Comparison of behavioral performance to ultimate limits and an ideal readout of the most sensitive ON-S and OFF-S RGC mosaics. The most sensitive OFF-S and ON-S RGCs were linked to behavioral performance on the decrement detection task using an ideal observer with access to the spike output from the full OFF-S or ON-S mosaic. The spike output was extracted using a three-stage process: (A) The tracked head position and direction let us determine the mouse’s relative position to the stimulus throughout a trial. (B) This relative position was used to determine the size and the location of the moving stimulus on the retinal mosaic. (C) Time-space separable linear-nonlinear Poisson (LNP) models, constrained from electrophysiology data collected in matching conditions, finally predicted the spike output of the full OFF-S or ON-S RGC mosaic. (D) Single-photon responses traverse the rod bipolar pathway. Ideal task performance in photon space (I) is degraded as losses and noise are introduced along the circuit, from rods (II) to RGCs (III). (E) Data points: behavioral performance (mean percentage of correct choices ± SEM, reproduced from Figure 3E). Dotted lines: ultimate detection limits computed in I) photon space, (IIa) observing the loss of photons not absorbed and not causing isomerizations in rods, (IIb) observing the same as in IIa and the effect of rod dark events (noise), and (III) observing the same as in IIb and the loss of single-photon responses in the rod-to-rod-bipolar synapse. Solid lines: ideal-observer readouts of the spike output of the full mosaic of OFF-S RGCs (dark blue) and ON-S RGCs (light blue).

We compared the ideal observer performance based on OFF-S and ON-S readouts to behavioral performance and to ultimate detection limits at different hierarchical levels of the rod bipolar pathway. These limits were estimated by a model (STAR Methods) that assumed pooling from an area matching the entire spatial extent of the stimulus and takes into account estimated photon losses and noise originating from spontaneous isomerizations of visual pigments in rods, i.e. pigment noise (0.01 R*/rod/sec; Burns et al. 2002) (see Figure 4D; I photon space, IIa rods, IIb rods + pigment noise, III RGCs + pigment noise). First, we found that behavioral performance exceeds that of the ideal readout of the ON-S RGCs, whereas there is a close match between behavior and the ideal readout of the OFF-S RGCs (Figure 4E: OFF model vs. Behavior). The ideal OFF-S RGC observer outperformed the ideal ON-S RGC observer by a factor of ~8 (Figure 4E: OFF vs. ON model). Second, while the performance of an ideal observer looking at the OFF-S RGC responses fell ~400-fold below the ultimate theoretical limit defined in photon space (Figure 4E: I vs. OFF model), it was only ~3-fold below the limit at the RGC level, where known retinal losses and noise were included (Figure 4E: III vs. OFF model). This result shows that OFF RGCs provide a remarkably good readout of their pooled rods, whereas both photon losses (Figure 4E: I vs. IIa, ~7-fold; IIb vs. III, ~2-fold: total ~14-fold) and retinal noise (Figure 4E: IIa vs. IIb, ~8-fold) fundamentally limit the detection of the weakest quantal shadows.

## DISCUSSION

Our results show that behavioral detection of the dimmest light decrements, here termed quantal shadows, relies on the most sensitive retinal OFF RGCs. The exquisite sensitivity of OFF-S alpha RGCs lets mice detect the absence of even a few photons in a pool of thousands of rods at starlight-level illumination. Our modeling shows that the main factors limiting the detection of quantal shadows are retinal losses of single-photon signals and retinal noise, thus suggesting an almost perfect readout of the signals from the OFF-S RGC mosaic in the rest of the brain to drive visually guided behavior.

Our results, together with the earlier results on light increment detection (Hecht et al., 1942; Smeds et al., 2019; reviewed in Kiani et al., 2020), show that the retina possesses fine-tuned circuit mechanisms for the detection of sparse photons (light increments) as well as decrements in extremely sparse photon fluxes. The shared need for high-fidelity processing of the weakest light increments and decrements is implemented by the rod bipolar pathway, where the split into ON and OFF pathways occurs at a late stage. This late split allows both pathways (ON and OFF) to utilize the large spatial pooling of rod signals required for high sensitivity. Intriguingly, this lets OFF-S RGCs encode decrements in dim backgrounds with a quantal sensitivity comparable to that of the most sensitive RGCs to light increments in darkness. The late ON and OFF split appears as an optimal tradeoff between distinct computational needs for the two channels and the shared need for high rod convergence. Considering that ON and OFF pathways represent a common theme across sensory modalities in the central nervous system (Chalasani et al., 2007; Gallio et al., 2011; Scholl et al., 2010), our results suggest a general neural processing principle for ON and OFF pathways in situations where signals are extremely sparse: A late split gives less space for computational divergence but better signal-to-noise ratios via additional pooling. In the case of the retina, the sparseness of signals sets tight demands for rod convergence and fundamentally limits how early the split can happen. Indeed, at higher photopic light levels, where photons are abundant, cone signals diverge into ON and OFF signaling routes already at the first synapse, thus favoring computational diversity rather than sensitivity via multiple distinct ON and OFF bipolar channels (Yan et al., 2020) feeding into >40 distinct ON and/or OFF RGC types (Baden et al., 2016; Bae et al., 2018; Rheaume et al., 2018).

Secondly, our results show that behavioral performance in detecting quantal shadows gets very close to an optimal readout of the most sensitive OFF-S but not ON-S RGCs. Together with our recent findings showing that the most sensitive ON RGCs, but not OFF RGCs, mediate detection of the weakest light increments in mice (Smeds et al., 2019) and in humans (Kilpeläinen et al., 2021), this shows that there is a categorical division of labor between ON and OFF pathways already at starlight levels in encoding lights and shadows for guiding behavior. These results further indicate that starlight-level behavioral tests can assess directly and with ganglion-cell type-specificity the functioning of retinal circuits – a feat that is incredibly difficult to accomplish at high light levels where behavior relies on >40 distinct RGC types. We suggest that single-photon and quantal-shadow behavioral detection paradigms could offer a unique avenue for probing end-to-end and cell-type-specifically ON and OFF channel function in healthy and diseased visual systems.

## ACKNOWLEDGEMENTS

We thank Drs. Kristian Donner, Greg Schwartz, Gabriel Peinado and Christophe Ribelayga for valuable comments on the manuscript; Matthew Dunkerley and Sathish Narayanan for the design of the data acquisition software; Dr. Martta Viljanen for technical assistance with the design of the water maze and Mr. Sami Minkkinen for technical assistance with the design of the patch rig. Support was provided by the Academy of Finland (296269, 305834 to P.A.-L.); the Aalto Brain Centre (J.W.); Svenska kulturfonden (J.W.); The Finnish Society of Sciences and Letters (J.W.); the European Union’s Horizon 2020 research and innovation programme (Marie Sklodowska-Curie grant 713645 to N.M.); the Doctoral programme Brain & Mind, University of Helsinki (S.K.); the Finnish Cultural Foundation (S.K.); the Ella and Georg Ehrnrooth Foundation (T.T.); the Oskar Öflund Foundation (T.T.); the Finnish Foundation for Technology Promotion (T.T.). We acknowledge the computational resources provided by the Aalto Science-IT Project.

## AUTHOR CONTRIBUTIONS

J.W., S.K., and P.A-L. designed the experiments. T.T. developed the methodology for markerless mouse tracking. T.T. and S.K. set up the water maze used for behavioral experiments and S.P carried out behavioral experiments. S.K and S.P collected and analyzed pupil data. N.M carried out RGC experiments. J.W, S.K and S.P analyzed data. J.W., S.K. and P.A-L wrote the MS.

## STAR METHODS

### RESOURCE AVAILABILITY

#### Lead contact, materials, data and code availability

Requests for further information or resources and reagents should be directed to and will be fulfilled by the Lead Contact, Petri Ala-Laurila. The data and custom code that support the findings from this study are available from the lead contact upon request.

## EXPERIMENTAL MODEL AND SUBJECT DETAILS

### Mice

Melatonin proficient mice (CBA/CaJ; Jackson Laboratory, Bar Harbor, ME, USA; males and females at the age of 8–21 weeks) were used in this study following similar procedures as described earlier (Koskela et al., 2020). Mice were housed in a 12 h/12 h light/dark cycle (white light ~300 lux; lights on from 20:00 to 8:00 in the animal housing room). The mice were dark-adapted for a minimum of 2–3 hours before all experiments. All animal procedures were performed according to the protocols approved by the Regional State Administration Agency for Southern Finland.

## METHODS DETAILS

### Ganglion cell experiments

All ganglion cell recordings were done from flat-mount preparations as described in earlier publications (Smeds et al., 2019; Koskela et al., 2020). Briefly, dark-adapted mice were sacrificed by rapid cervical dislocation, their eyes were enucleated, hemisected, and the vitreous was removed. The eye cups were stored in a light-tight container at 32°C in oxygenated (95% O_2_/5% CO_2_) Ames solution (Sigma, A-1420; osmolality adjusted to 280 ± 2 mOsm/kg). The whole retina was gently isolated from the pigment epithelium and placed on a poly-D-lysine coverslip (12-mm; VWR, Corning) with the photoreceptor side down. The mounted retina was then placed on a recording chamber and transferred to the microscope (SliceScope Pro 3000, Scientifica) and perfused with Ames solution (32 ± 1°C, flow rate: 8 ml/min). These and all subsequent procedures were performed under I.R. light (>900 nm) using night vision goggles (PVS-7-1600, B.E. Meyers) and I.R. pocket scopes (D7200-I-1600, B.E. Meyers) attached to the dissection microscope. The electrodes for RGC recordings (4–6 MΩ) were filled with Ames. The preparations were visualized using I.R. light (940 nm; turned off during recordings) and a CCD camera (Wat-902HS, Watec) attached to the microscope. Experiments were done at the subjective day of the mice (absolute sensitivity in darkness is not affected by the diurnal rhythm; Koskela et al., 2020).

We targeted ON-S and OFF-S alpha RGCs and carried out spike recordings using the cell-attached patch-clamp technique on flat-mount retina preparations. These RGC types were identified based on the large soma size and their characteristic light response. In a subset of experiments, the dendritic morphology of cells was verified by filling the cells with a fluorescent dye (HiLyte Fluor 750 hydrazide, AnaSpec, AS-81268) and by imaging the cells (Andor Zyla 4.2 PLUS sCMOS) following fluorescence excitation (peak at 740 nm; width 35 nm, CoolLED pE-4000, CoolLED). The cell morphology was confirmed to be consistent with ON-S and OFF-S alpha RGCs (Smeds et al., 2019). The quality (high sensitivity) of the mount was ascertained by verifying that all RGCs had 2AFC thresholds below 0.05 R*/rod in darkness (Figures 2I and 2J) and that the baseline firing rate of ON-S RGCs was below 5 spikes/s.

#### Light stimulation (RGC)

Calibrated visual stimuli centered on the target cell were created with a DLP projector (912 × 1140 pixels; 1.8 μm/pixel on the retinal surface; 60 Hz frame rate; blue LED spectral peak ~450 nm; Texas Instruments, LightCrafter 4500). The stimulus was focused on the photoreceptor plane by the microscope condenser localized beneath the preparation. The flat-mounted retina (photoreceptor downwards) was thus stimulated directly from the photoreceptor side. The stimuli were circular spatially uniform (340 μm in diameter) increments (in darkness) or decrements (340 μm in diameter dark spot) from a square spatially uniform background (1,000 × 1,000 μm). The increments were always short flashes in darkness (17 ms) used to validate the sensitivity of the cell, whereas various stimulus durations (17–400 ms) were used for light decrements across dim background lights (darkness–30 R*/rod/s). The decrement stimuli were always as brief as possible, meaning that durations longer than 17 ms were only used with maximal contrast. The intensity was set by adjusting the LED current in the projector and by inserting calibrated neutral density filters (Thorlabs) into the light path. The cells were always given time to adapt when changing the background light (2 min. at background lights < 1 R*/rod/s and 5 min. at background lights > 1 R*/rod/s).

### Behavioral experiments

Behavioral experiments were carried out in a homogeneously lit white polyethene six-armed water maze in dim background light to monitor the ability of dark-adapted mice to find a dark stimulus spot across a range of dim background lights. The physical maze was inspired by previously used water mazes (Hayes and Balkema, 1993; Sampath et al., 2005) but modified for our use by using a smaller black stimulus spot and making the setup compatible for markerless tracking of mouse swimming trials. The maze was surrounded by white satin curtains, and non-fat (skim) milk was added to the water (depth: 8 cm; temperature: ~20°C) to make it opaque and the escape ramp invisible for the mice. The stimulus was a non-reflecting black spot mounted on a movable white polyethene wall in one of the six arms of the maze. The location of the stimulus across the six arms was randomized across trials. The stimulus contrast was measured to be ~97% (Weber), and the stimulus diameter was 4 cm (~170 μm in diameter when projected to a mouse retina located at the center of the maze).

The visual sensitivity of mice was determined as previously described (Smeds et al., 2019; Koskela et al., 2020). Briefly, a mouse was placed in the center of the maze in a transparent tube and allowed to orient itself for ~5 s. The tube was then removed, and the mouse was free to search for the stimulus located in one of the six arms of the maze. A choice was defined as correct if the mouse entered the stimulus corridor as its first choice. The mice (4 females, 6 males) were initially trained to associate the escape ramp with the stimulus. Training was done with a constant bright background light (~10,000 photons/s/μm^2^) and continued until the mice reliably made a correct choice in ≥ 80% of the trials (this took 15 days). Subsequently, the experiment phase began, and the background illumination was attenuated (testing one intensity per day) until the mice made a choice completely randomly. After the experiment phase, the mice were re-tested using a high background intensity to ensure that no significant changes had occurred in their overall ability to perform the task as well as complete darkness to ensure that they did completely random choices and there were no biases or a priori probabilities for favoring particular arms of the maze (Figure 3A). The order of the mice (males or females first) was alternated every second day. Each mouse performed four trials per day during all phases (training, experiment, and control). The experiments were done during the subjective night of the mice (3h from light offset, as described in Koskela et al., 2020). Every trial was recorded using I.R. light (Osram OSLON Black Series 940 nm high-power LEDs) and a sensitive CMOS camera (Basler ace acA1920-50GM with Basler Pylon viewer 64-bit software).

#### Light stimulation (behavior)

The white maze was illuminated by an LED (peak at 515 nm, Cyan Rebel LED, Quadica MR-E-0077020S) connected to a lens tube with a bandpass filter (λ_max_ ~510 nm, FWHM ~10 nm, Thorlabs) and neutral density filters (Thorlabs). The light path was then split at the end of the lens tube using a trifurcating fiber (Newport). The three fiber ends were equipped with convex lenses (N-BK7 plano-convex, Thorlabs), placed in a symmetric arrangement around the maze, and directed to spread the light onto a photographic reflector (Lastolite L3831, diam. 95 cm) located above the maze. This yielded a homogeneous illumination over the maze, and the homogeneity was checked before, during, and after the full experiment (see light calibrations below). The light intensity was set by using suitable neutral density filters and by controlling the current driving the LED, and carefully calibrated (see light calibrations below).

#### Automated tracking and stimulus projections

The body position and head direction of mice were automatically and markerlessly tracked during each trial. The tracking system will be explained in details in a separate technical paper (T.T. and P.A.-L., unpublished data), but briefly, it consisted of 1) a deep convolutional neural network that detected keypoints, 2) a state-space model that fused the keypoints into an estimated head position and direction, and 3) a smoother that filtered the raw state estimates to handle occlusions and remove noise. Stimulus projections onto the retinas across the swimming trials were estimated as in Smeds et al. (2019) by utilizing typical values for eye sizes and their anatomical locations. Compensatory eye movements were estimated based on previous measurements (Smeds et al., 2019).

### Light calibrations

#### RGC recordings

Light intensities were calibrated with an optometer (S470 with the sensor 268R, UDT Instruments), and the irradiance spectrum was measured with a spectrometer (Jaz spectrometer, OceanOptics). The calibrated photon fluxes were converted to photoisomerization rates per rod (R*/rod/s) using the measured LED spectrum, the absorption spectrum for rhodopsin (peak sensitivity at 497 nm: Govardovskii et al., 2000; Toda et al., 1999), and a rod collecting area of 0.6 μm^2^ (Smeds et al., 2019; Koskela et al., 2020).

#### Behavioral experiments

Light intensities were calibrated with an optometer (S470 with the sensor 268R, UDT Instruments) and a luminance meter (Konica Minolta LS-110). The spectral irradiances of the stimuli were measured with a spectrometer (Jaz spectrometer, OceanOptics). The corneal flux densities, given in photons/s/μm2, were obtained from optometer recordings at the center of the maze, with the sensor oriented to collect light along the horizontal axis from one of the maze arms. The power (P_center_) recorded in this way was converted to a corneal photon flux density (F_cornea_) as:

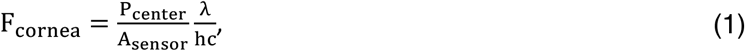

where λ denotes the peak of the bandpass filter (510 nm), h is Planck’s constant, c, is the speed of light, and A_sensor_ is the sensor area (1 cm^2^ in our case).

To convert the flux density to isomerization rates per rod (R*/rod/s) requires the amount of reflected light per wall area to be quantified. This was done by repeating the optometer measurements at the center of the maze, as above, with and without the near-perfectly dark stimulus located at the back of the maze arm. We further mounted a 3D-printed cone on top of the sensor to restrict the collecting angle and thus increase the relative difference. The difference in the measured power (P_diff_) thus represented the amount of light reaching a retinal area of the same size as the projected stimulus (A_retina_: 22,700 μm^2^, see Smeds et al., 2019). The intensity (I) in isomerizations per rod per sec could therefore be determined as:

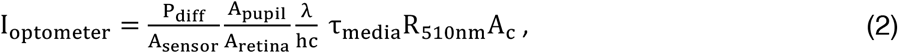

where A_pupil_ was stable over the relevant backgrounds (mean 6.5 mm^2^, see pupil measurements below), τ_media_ is the light transmittance of the ocular media (0.55; Henriksson et al., 2010), R_510nm_ is the relative absorption factor of rhodopsin at 510 nm (0.93 for a peak sensitivity at 497 nm; Govardovskii et al., 2000; Toda et al., 1999), and A_c_ is the rod collecting area (0.6; Smeds et al., 2019). As a control, we also determined the isomerization rate based on luminance measurements, as the luminance meter quantifies the amount of light reflected per area directly. The measured luminance (L; photopic) was converted to an isomerization rate as:

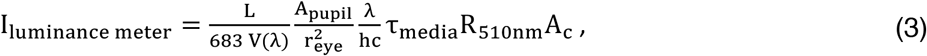

where V(λ) is the photopic luminosity function, and r_eye_ is the eye radius (1.7 mm; Remtulla and Hallett, 1985). Both methods yielded the same results (within measurement accuracy).

The uniformity of the illumination was verified regularly by measuring both the corneal flux density and the corresponding isomerization rates for each corridor. The maximal difference remained below 10% (from the mean) throughout the experiment.

### Pupil measurements

The pupil areas (A_pupil_, see above) of mice were measured in darkness and across dim backgrounds (in matching conditions with behavioral experiments). These measurements allowed us to estimate the light intensities used during behavioral experiments as photoisomerizations rates in rods (R*/rod/s). The procedure is described in detail in Smeds et al. (2019). Briefly, the I.R. LEDs of the experimental setup were turned on to make the pupils discernable for imaging using an I.R.-sensitive camera (Basler ace acA1920-155um). The mouse was held by its tail in the dry water maze using otherwise identical conditions with the behavioral measurements. The pupil areas were then measured from single frames in the recorded video by tracing out the border of the pupil in each frame using ImageJ (1.47v, National Institute of Health, USA).

### Data analysis

#### Sensitivity of RGCs

The RGC response was quantified as the average spike count difference (between post-stimulus and pre-stimulus firing rates). The time window and the sign of the response (spike increase or decrease) were determined based on the average response of the cell at each background. The sensitivity, or response per absorbed photon (spikes per R*), was obtained by dividing the difference by the stimulus magnitude and by multiplying by the rod convergence per RGC (~10 000; Dunn et al., 2006; Tsukamoto et al., 2001). Normalized sensitivities were obtained by dividing with the sensitivity in darkness. The fitted gain (G) functions in Fig. 2 E and G are generalized Weber functions:

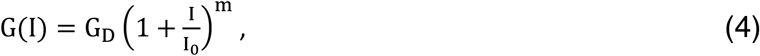

where I is the background, G_D_is the gain in darkness, I_o_ determines the onset of adaptation, and m is the slope. The parameters were always fitted to data using a least squares objective function.

#### Discrimination performance of RGCs

Two-alternative forced choice (2AFC) ideal observer analysis was performed to determine the sensitivity threshold of RGCs for increments (in darkness) and decrements across dim backgrounds (Ala-Laurila and Rieke, 2014; Chichilnisky and Rieke, 2005; Smeds et al., 2019). Briefly, the instantaneous firing rate was computed for 20 ms time bins, and a discriminator was created from the mean response over all trials at the same background intensity. The response magnitude for each trial was quantified by the dot product between the discriminator and the instantaneous firing rate during a 750 ms window starting at stimulus onset. Similarly, the noise magnitude was quantified in an identical manner but by using a 750 ms window that ended at stimulus onset. The percent correct rate for a 2AFC task at any stimulus intensity was evaluated by determining how often the response magnitude was larger than the noise magnitude within the same trial. The 75 % correct choice threshold was finally found through interpolation between the percent correct rates obtained at each stimulus magnitude. The complete procedure was then repeated a second time for decrements using ON-S RGCs, but this time using only negative values in the mean discriminator to evaluate the fraction of ON-S RGCs that could encode the stimulus using their onset response only. The gray numbers in Figure 2J thus correspond to the fraction of ON-S RGCs for which we obtained 75 % thresholds with the normal mean discriminator but not with the constrained discriminator that only looked at the onset response.

#### Psychometric function

The behavioral data (percent correct) was fit with a psychometric function of the form:

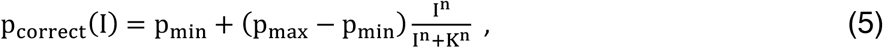

where p_min_ denotes chance level (1/6), p_max_ is the maximal performance (1), K is the intensity at ~0.58, and n is the slope. Optimal K and n parameter values were found by minimizing the least squared error between the measured data and the predicted values of the psychometric function.

#### Analysis of the ultimate detection limits

The ultimate limit for RGC performance was defined as ideal performance in distinguishing between two Poisson distributed variables, signal and noise response distributions, in a 2AFC setting. The mean of the signal distribution was set by the product of the rod convergence per ganglion cell (~10 000; Dunn et al., 2006; Tsukamoto et al., 2001), the spontaneous isomerization rate of rods (0.01 R*/rod/s; Burns et al., 2002; Fu et al., 2008), the probability that single-photon responses are transmitted from rods to rod-bipolar cells (0.25, Field and Rieke, 2002), and the duration of the time window over which isomerization events were counted. The time window corresponded to 50 ms (Field et al., 2019) plus the time needed for presenting a decrement of the required magnitude, thus giving rise to increasing thresholds for backgrounds dimmer than 0.01 R*/rod/s. The mean of the noise distribution was defined in an equivalent manner, but by also incorporating the isomerization rate due to the background light.

The ultimate limits for behavioral performance, in turn, were taken as ideal performance on a 6AFC task, where an ideal observer had to make a choice between six alternatives (one signal and five noise samples). The signal and noise samples always came from Poisson distributions, whose means (λ_noise_ and λ_signal_) were defined by the level of comparison. That is, for the photon level, we used λ_signal_ = 0, whereas λ_noise_ depended on the background and the duration (t) of the counting window. For the rod level, λ_signal_ was again zero, but this time λ_noise_ was scaled by ~0.14 due to losses in converting photons to isomerizations (see detailed description below). At the rod noise level, the contribution of spontaneous isomerizations (0.01 R*/rod/s; Burns et al., 2002) was added to both means (0.01N_rods_t), and at the RGC level, we additionally considered losses at the rod to rod-bipolar synapse, by further scaling both means by 0.25 (Field & Rieke, 2002). The number of rods N_rods_ was fixed at 20176 (based on the stimulus size of 40352 μm^2^ 10 cm from the center of the maze and a rod density of 500 000 rods/mm^2^; Volland et al., 2015), and the duration of the counting window was fixed at 320 ms (based on matching ideal observer performance to behavior in Smeds et al., 2019). The final conversion from photon fluxes to isomerization rates was obtained by scaling the photon flux with the scaling factor due to conversion losses (0.14, see below) divided by N_rods_.

#### Losses in converting photons to isomerizations

The efficiency (eff.) by which photons arriving at the mouse cornea are converted to isomerizations was determined by accounting for known losses (l) as:

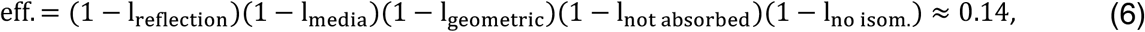

with l_reflection_ = 0.04 (Hecht et al., 1942), l_media_ = 1 − τ_media_ = 0.45 (Henriksson et al., 2010), l_geometric_ = 0.4 (based on total rod coverage and light funneling; Lyubarsky et al., 2004), 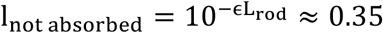 (ϵ = 0.019 *μ*m^−1^ and L_rod_ = 24 *μ*m; Lyubarsky et al., 2004), and l_no isom_ = 0.33 = 1 − Q (quantum efficiency, Baylor et al., 1979). In total, this resulted in a total loss of ~0.86, meaning that roughly 14% of all photons arriving at the mouse cornea resulted in an isomerization.

#### Ideal RGC observer

We used an ideal observer model for correlating the responses of populations of ON-S or OFF-S RGCs with behavioral data (following similar procedures as in Smeds et al. 2019). This ideal observer had access to the modeled output from the full mosaic of either OFF-S or ON-S RGCs during behavioral trials. Each RGC in the mosaic was modeled using an LNP model (see below) tightly constrained by electrophysiology data collected in matching stimulus conditions. Ideal performance on the behavioral task was defined based on the same 6AFC task as was used for the ultimate limits above. The difference being that λ_signal_ was now defined as the total spike count from the 10 RGCs among all RGCs in the full mosaic giving the strongest response during the duration of the counting window (320 ms; as found in Smeds et al., 2019) while the mouse was within 10 cm of the center of the maze. This number of RGCs corresponds roughly to the number of ON-S or OFF-S receptive fields partially covered by the real visual stimulus in the two retinas of the mouse (see Smeds et al. 2019 for further details). Similarly, λ_noise_ was defined as the total spike count from 10 RGCs during the duration of the counting window when the RGCs fired at their background-dependent baseline firing rate (see Smeds et al., 2019 for a detailed descriptions of these assumptions). The sensitivity of the modeled RGCs was adjusted based on their 2AFC thresholds in darkness. We set the modeled OFF-S and ON-S RGCs to have a threshold that corresponded to the most sensitive RGCs recorded (OFF-S: 0.0013 R*/rod and ON-S: 0.0061 R*/rod; in line with the average ON-S threshold being ~4 times higher than for OFF-S RGCs).

#### LNP model

We used LNP models to predict ON-S and OFF-S RGC responses to visual stimuli in matching conditions with behavioral experiments. We represented ON-S and OFF-S RGCs with a time-space separable LNP model following a similar approach as in Smeds et al. (2019). The LNP model was equipped with additional gain and baseline functions (constrained by data) in addition to the standard temporal and spatial filters to account for changes across background intensities (Figure 4C). The nonlinearity was fixed and resembled a rectified linear function defined as:

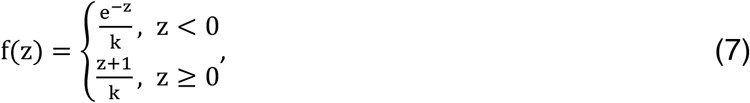

where the similarity score z constituted the sum of a bias term (for setting the baseline firing rate) and the cross-correlation between the scaled (background dependent gain) model filters and the stimulus. The parameter k was included for shifting the location of the split in the nonlinearity with respect to the baseline firing rate. The temporal filters as well as the parameter k were estimated from the stimulus response data obtained from all backgrounds (darkness, 0.03, 0.3, 3, and 30 R*/rod/s) by minimizing the negative log-likelihood function for a time-gain-separable LNP model (assuming that a fixed temporal filter is scaled by a gain factor at each background):

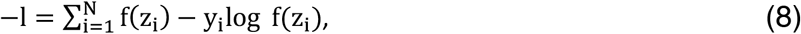

where y_i_ denotes the spike count within bin i. The time-gain separability constraint makes the log-likelihood function non-convex, and consequently, it was solved using an approach that solved two convex subproblems in alternation: Finding the temporal filter is a convex problem with the gain parameters fixed, and vice versa. The spatial filters, in turn, were taken to be Gaussian-shaped ( σ = 75 μm; Smeds et al., 2019), and the gain as well as the baseline functions were obtained from the data in Figure 2. The gain function for both ON and OFF RGCs was nonetheless obtained from the fit to the measured OFF RGC data, as this fit matched data from the ON RGCs over all backgrounds where we observed responses from the ON RGCs.

### QUANTIFICATION AND STATISTICAL ANALYSIS

All data analysis and result figures were done in MATLAB (version R2017B and later). All data are presented as mean ± standard error of mean (SEM) unless otherwise stated in the figure legends. The absolute thresholds for RGCs are summarized with the geometric mean and SEM. Details of the sample size (n) for each experiment can be found in the figure and/or in the figure legend. The statistical significance of the differences in sensitivity thresholds was tested using Welch’s t-test, whereas a paired t-test was used to assess the difference in pupil size between darkness and a background of 1 R*/rod/s. The normality of the data was tested using the Kolmogorov-Smirnov normality test. A p-value of 0.05 was used to define significance in all test. All tests used for p values were two-tailed.

## Notes

### Competing Interest Statement

The authors have declared no competing interest.

